## Supplementary figures and images for "Intermittent Bulk Release of Human Cytomegalovirus"

### Supplementary Figure 1.tiff

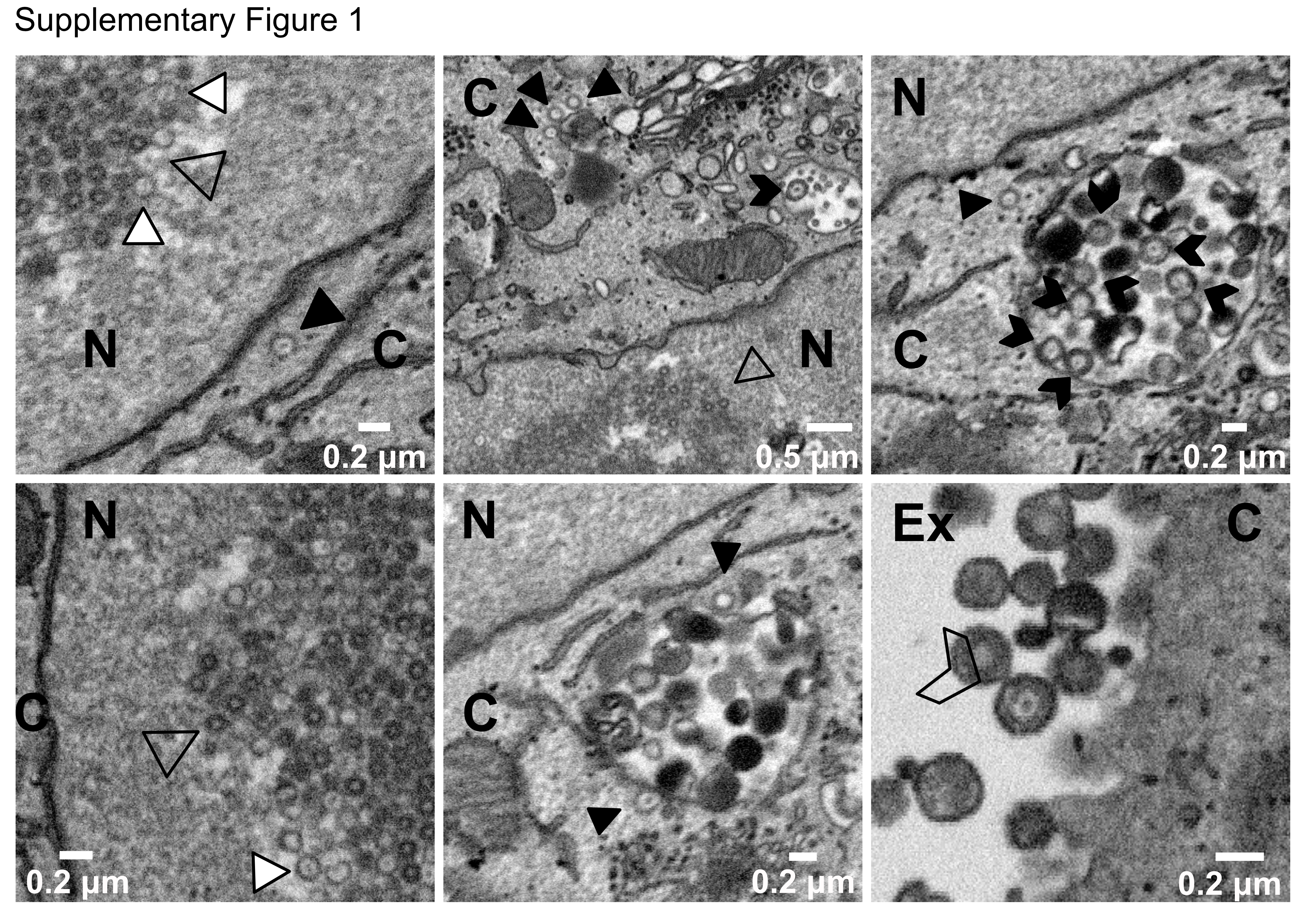

### Supplementary Figure 2.tiff

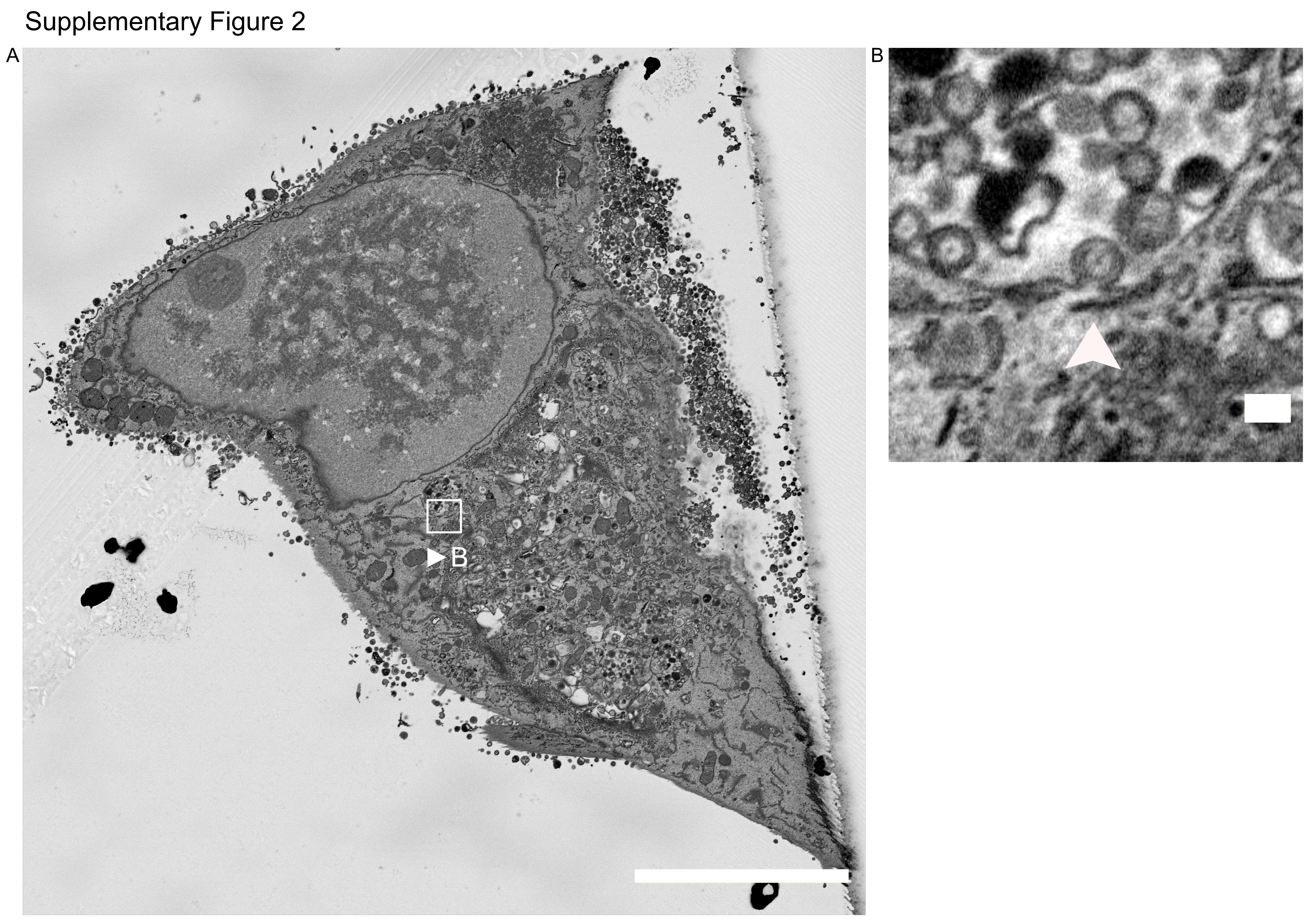

### Supplementary Figure 3.tiff

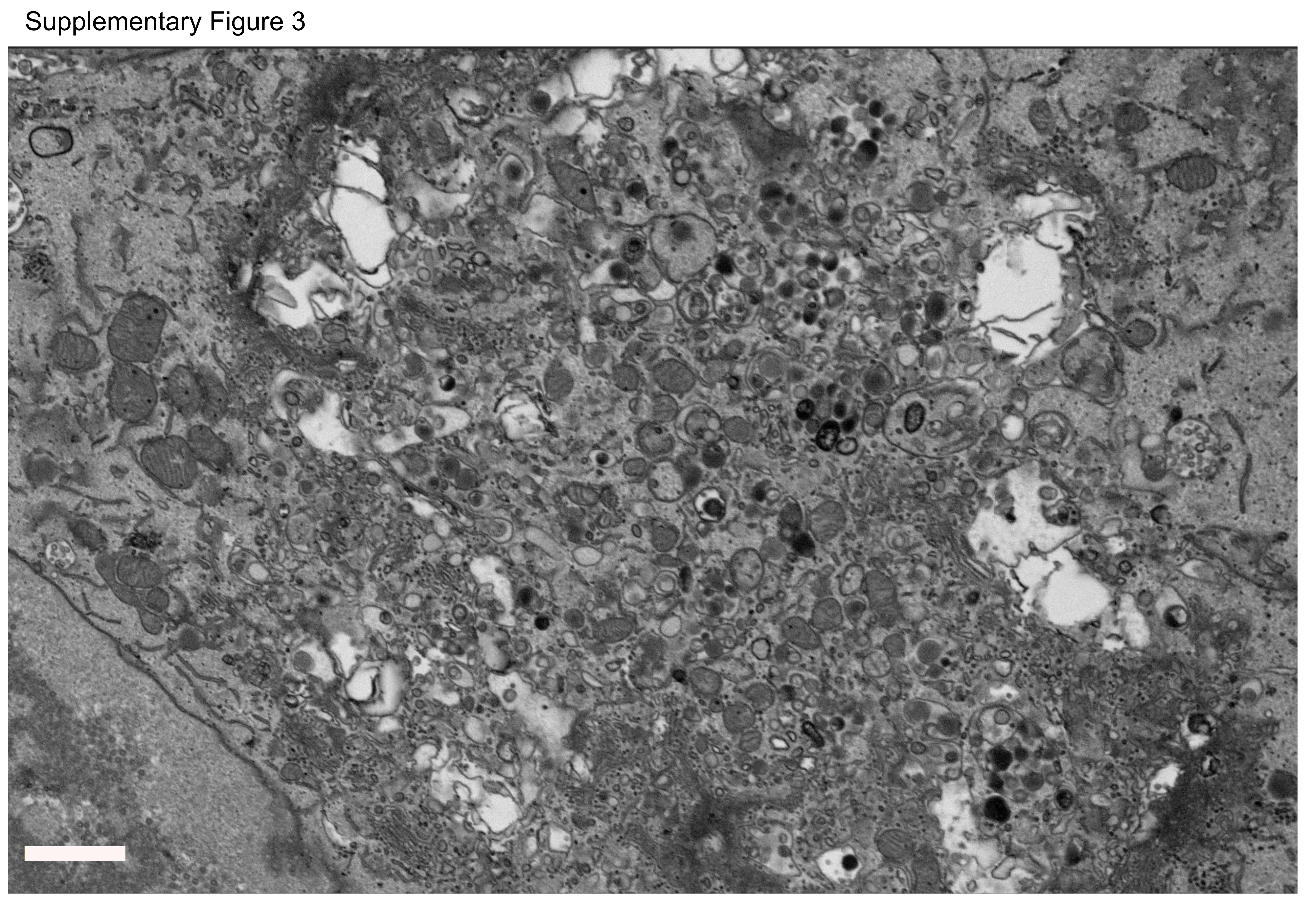

### Supplementary Figure 4.tiff

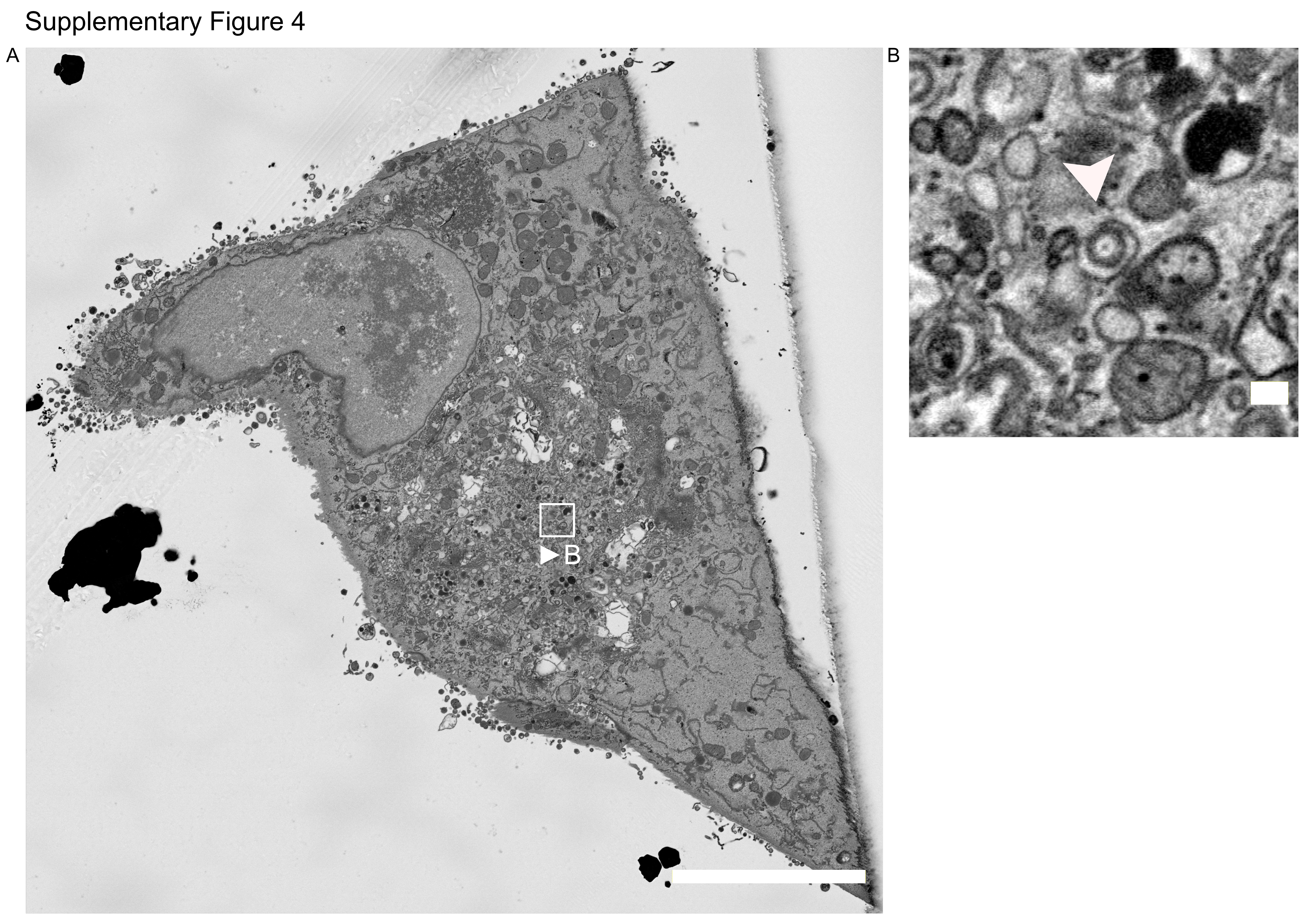

### Supplementary Figure 5.tiff

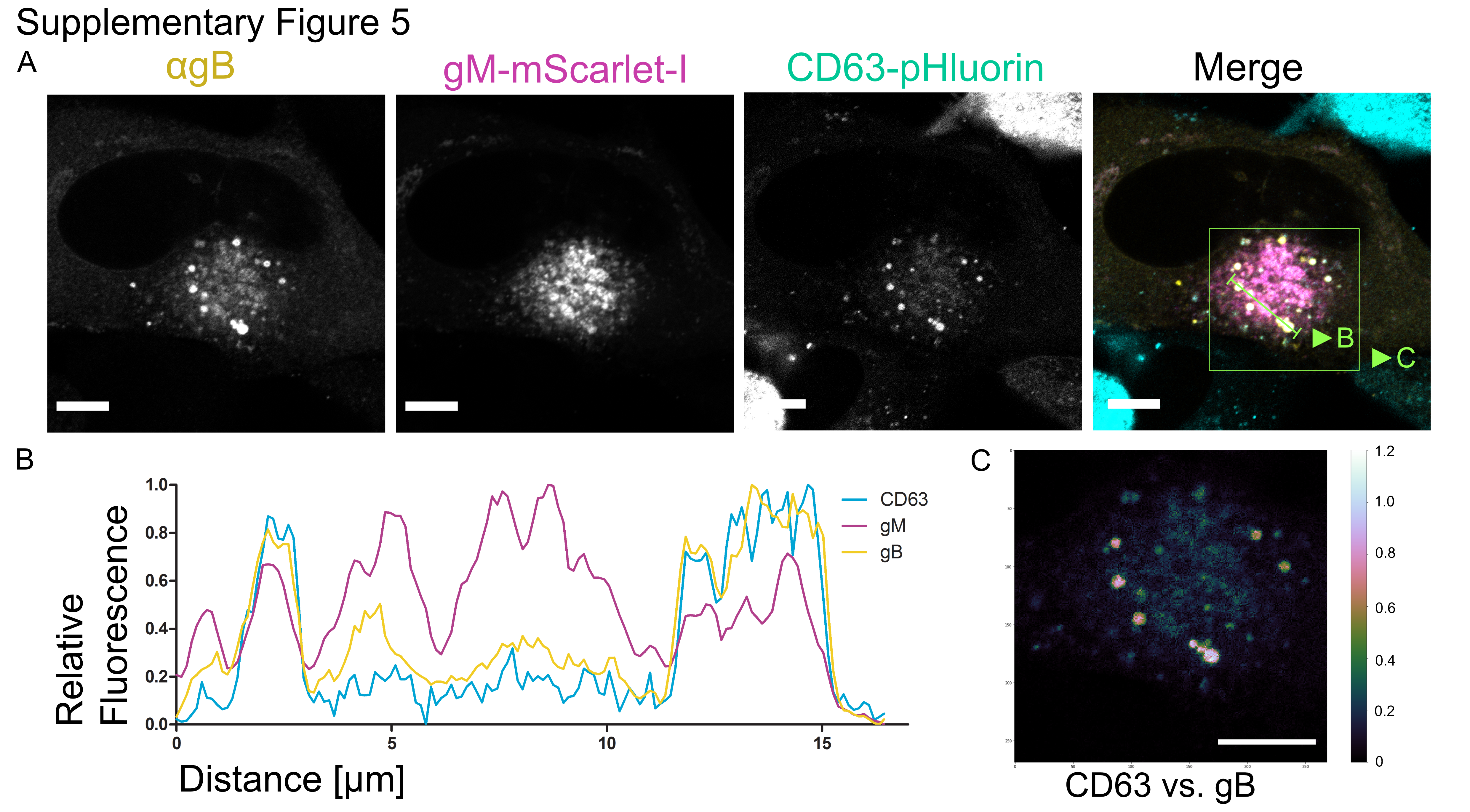

### Supplementary Figure 6.tiff

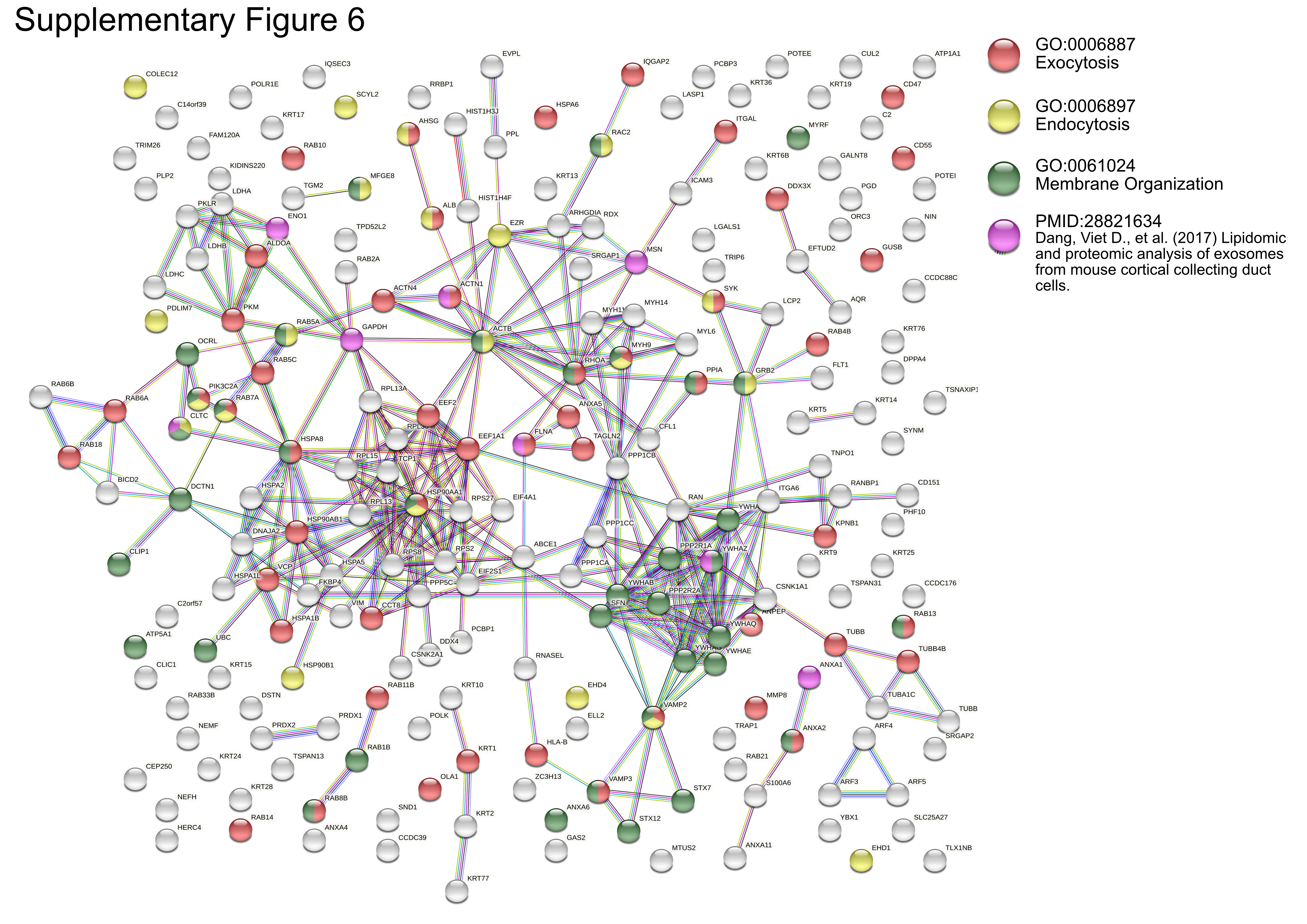

### Supplementary Figure 7.tiff

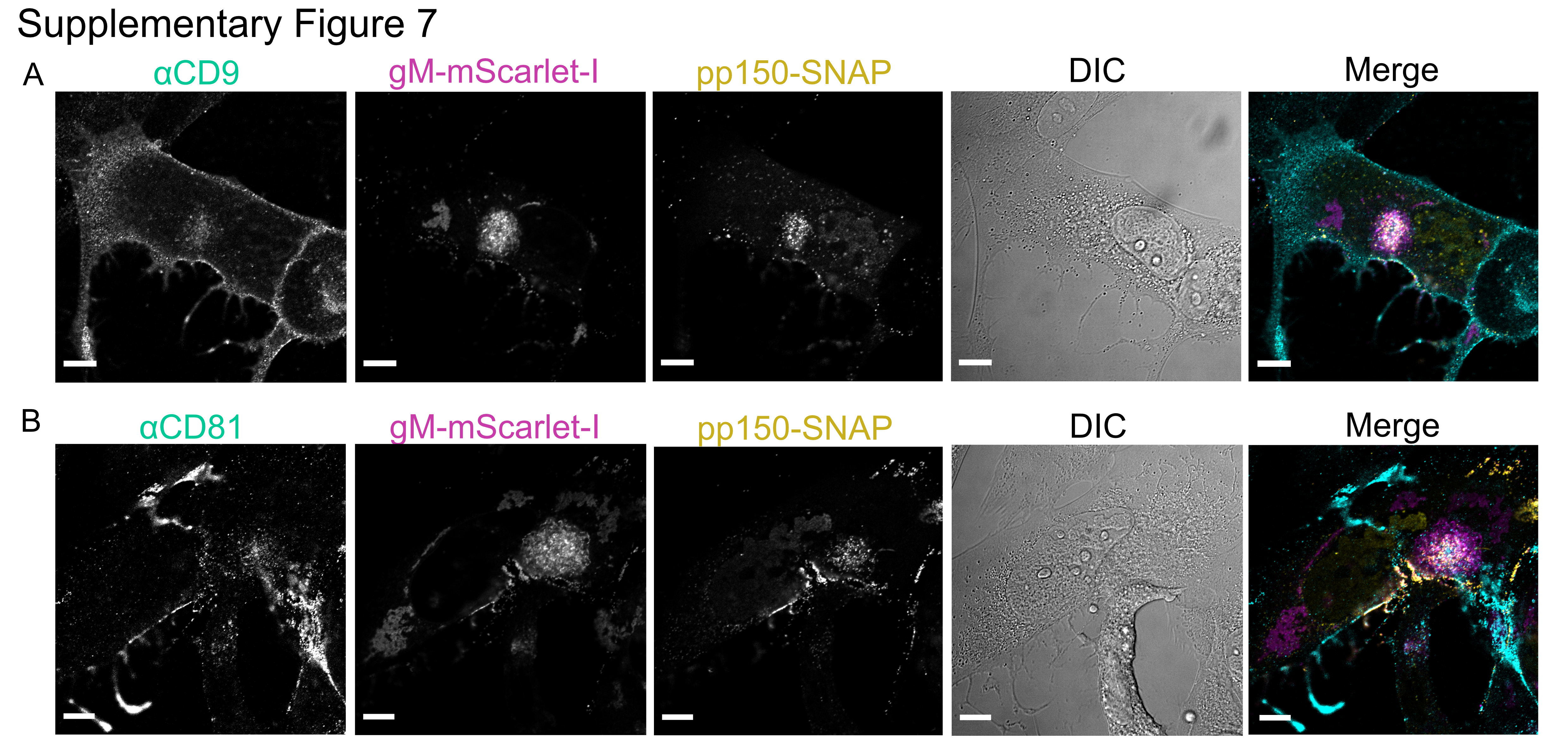

### Supplementary Figure 8.tiff

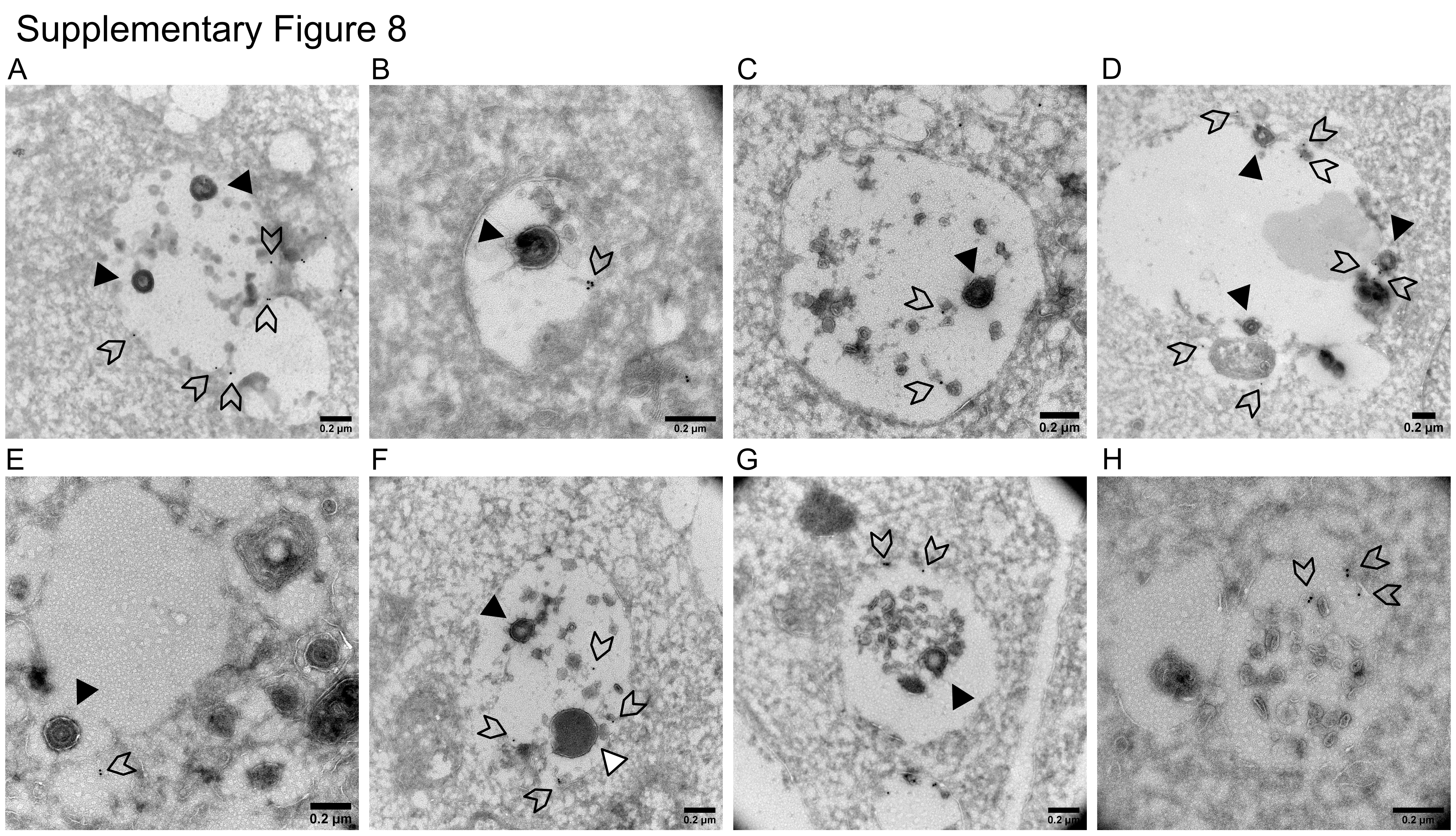

### Supplementary Figure 9.tiff

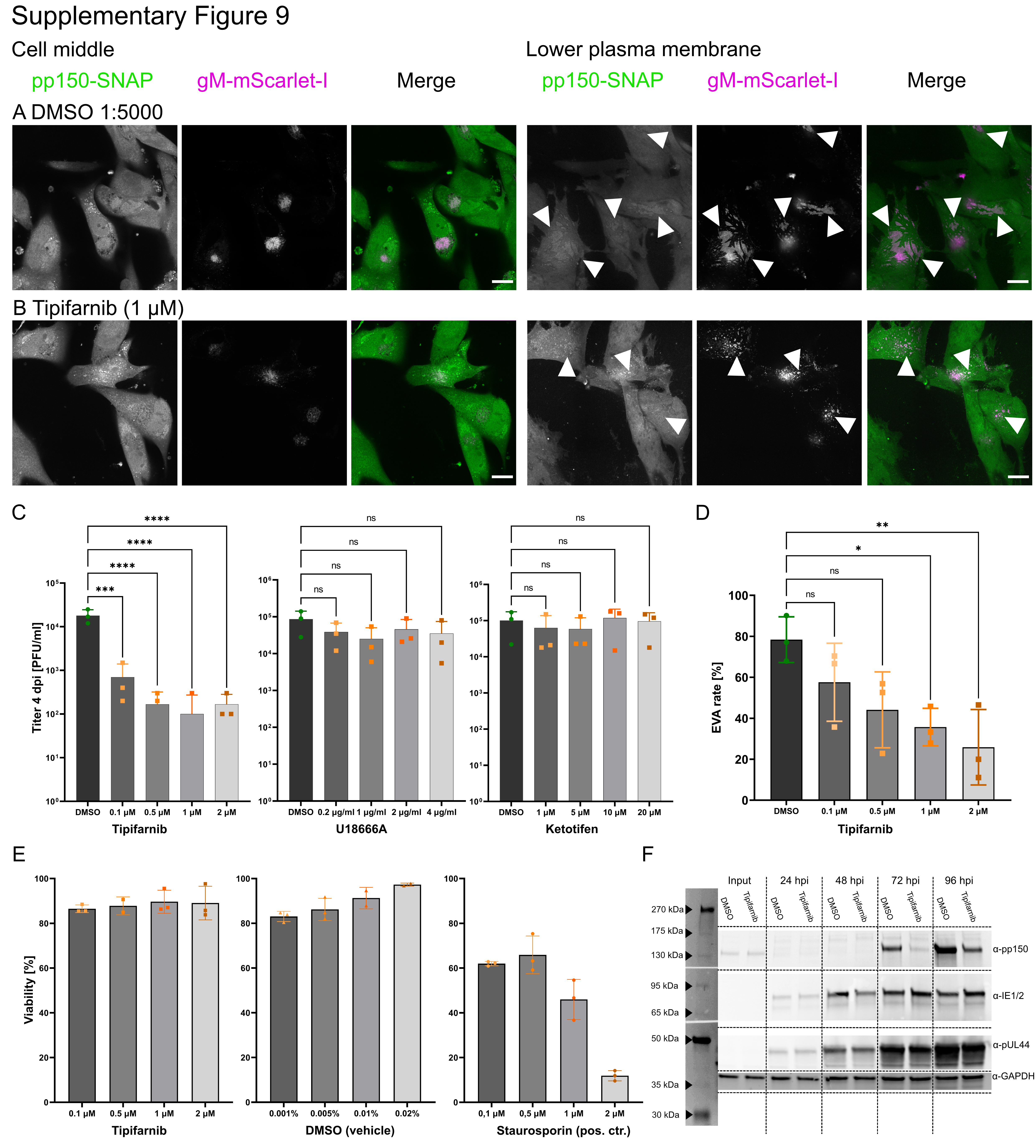
